# Evaluating state space discovery by persistent cohomology in the spatial representation system

**DOI:** 10.1101/2020.10.06.328773

**Authors:** Louis Kang, Boyan Xu, Dmitriy Morozov

**Affiliations:** Redwood Center for Theoretical Neuroscience, University of California, Berkeley; Neural Circuits and Computations Unit, RIKEN Center for Brain Science; Department of Mathematics, University of California, Berkeley; Computational Research Division, Lawrence Berkeley National Laboratory

## Abstract

Persistent cohomology is a powerful technique for discovering topological structure in data. Strategies for its use in neuroscience are still undergoing development. We comprehensively and rigorously assess its performance in simulated neural recordings of the brain’s spatial representation system. Grid, head direction, and conjunctive cell populations each span low-dimensional topological structures embedded in high-dimensional neural activity space. We evaluate the ability for persistent cohomology to discover these structures for different dataset dimensions, variations in spatial tuning, and forms of noise. We quantify its ability to decode simulated animal trajectories contained within these topological structures. We also identify regimes under which mixtures of populations form product topologies that can be detected. Our results reveal how dataset parameters affect the success of topological discovery and suggest principles for applying persistent cohomology, as well as persistent homology, to experimental neural recordings.

## 1 Introduction

The enormous number of neurons that constitute brain circuits must coordinate their firing to operate effectively. This organization often constrains neural activity to low-dimensional manifolds, which are embedded in the high-dimensional phase space of all possible activity patterns [1, 2, 3, 4]. In certain cases, these low-dimensional manifolds exhibit nontrivial topological structure [5]. This structure may be imposed externally by inputs that are periodic in nature, such as the orientation of a visual stimulus or the direction of an animal’s head. It may also be generated internally by the network itself; for example, the grid cell network constructs periodic representations of physical space which outperform non-periodic representations in several ways [6, 7, 8, 9, 10, 11, 12]. In either case, detecting and interpreting topological structure in neural data would provide insight into how the brain encodes information and performs computations.

One promising method for discovering topological features in data is persistent cohomology [13, 14, 15]. By tracking how the shape of the data changes as we examine it across different scales—thickening data points by growing balls around them—persistent cohomology detects prominent topological features in the data, such as loops and voids. This knowledge helps to identify the low-dimensional manifolds sampled by the data, and in particular to distinguish between tori of different intrinsic dimensions. Furthermore, it enables parametrization of the data and navigation of the underlying manifolds.

We characterize how persistent cohomology can discover topological structure in neural data through simulations of the brain’s spatial representation system. This system contains several neural populations whose activity exhibits nontrivial topology, which we term *periodic neural populations* (Fig. 1A). Grid cells fire when an animal reaches certain locations in its environment that form a triangular lattice in space [16]. In each animal, grid cells are partitioned into 4–10 modules [17]. Within each module, grid cells share the same scale and orientation but their lattices have different spatial offsets. Modules appear to increase in scale by a constant ratio and exhibit differences in orientation [17, 18]. Head direction cells fire when an animal’s head is oriented in a certain direction relative to its environment [19]. They respond independently of the animal’s position. Finally, conjunctive grid × head direction cells respond when an animal is located at the vertices of a triangular lattice and is oriented in a certain direction [20]. Like grid cells, conjunctive cells are also believed to be partitioned into modules.

**Figure 1:**
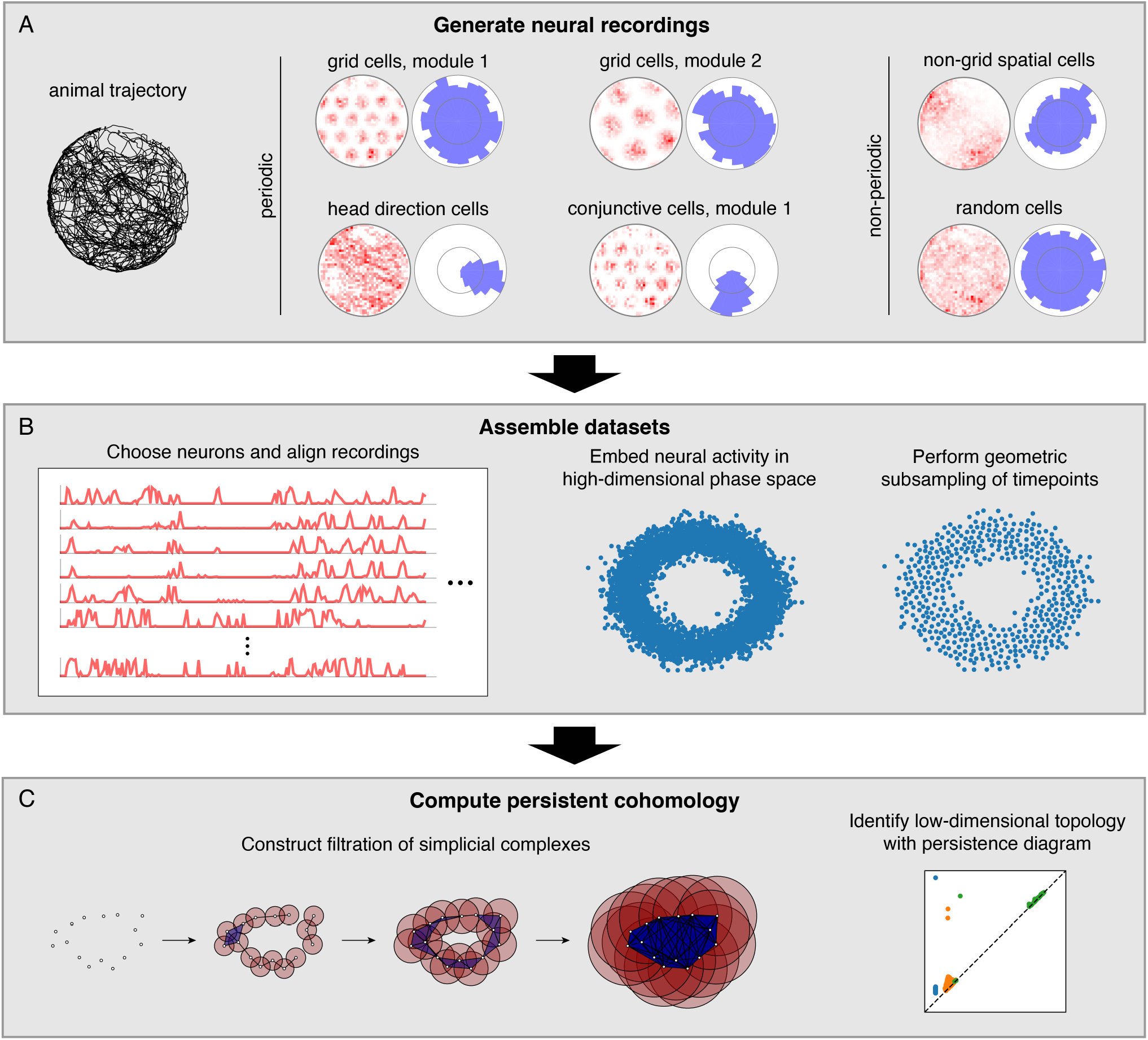
Pipeline for simulations and data analysis. (**A**) We generate activities for multiple neural populations along an experimentally recorded rat trajectory. For each population, we plot activity maps as a function of position (left) and direction (right) for one example neuron. (**B**) Then we choose neurons for topological analysis and form a high-dimensional vector of their firing rates at each timepoint along the trajectory. For computational tractability, we eliminate the most redundant points using a geometric subsampling algorithm. (**C**) We compute persistent cohomology on these subsampled timepoints to identify low-dimensional topological structure.

We also consider neural populations whose activity exhibits trivial topology, which we will term *non-periodic neural populations* (Fig. 1A). Place and non-grid spatial cells are part of the spatial representation system, and they fire in one or multiple regions of the environment [21, 22, 23]. These two populations are found in different brain regions, and the former tend to have sharper spatial selectivity compared to the latter. Finally, we simulate neurons with irregular activity that exhibits no spatial tuning. We imagine these *random cells* may be responding to non-spatial stimuli or representing internal brain states

Persistent cohomology, as well as the closely related technique persistent homology, has recently been applied to experimental neural recordings within the spatial representation system. It was used to discover topological structure [24, 25] and decode behavioral variables [24] from head direction cells. It is currently being implemented to do the same for grid cell recordings [26, 27], and researchers have demonstrated topological discovery in simulated grid cell data [25]. These works have improved our understanding of the large-scale organization of spatial representation circuits through persistent cohomology.

In contrast to the research described above, we aim to comprehensively explore the capabilities of persistent cohomology for simulated datasets. With complete control over the data, we can identify features that improve topological discovery and features that disrupt it. We can also freely generate datasets with varied quantities and proportions of different neural populations. A greater number of neurons embeds underlying activity manifolds in higher dimensions, which can strengthen the signal. However, experimental limitations impose bounds to this number. Our simulations allow us to evaluate persistent cohomology in regimes currently accessible by experiments, as well as in regimes that may soon become experimentally tractable due to advances in recording technology [28].

## 2 Results

### 2.1 Overview of methods and persistence diagrams

In this work, we simulate neural populations within the spatial representation system, prepare the simulated data for topological analysis, and compute persistent cohomology to discover topological structure within the data (Fig. 1). We will now briefly describe each of these three stages; a complete explanation is provided in the Methods section.

To generate neural recordings, we define tuning curves as a function of position and direction. For each grid module, we first create a triangular lattice in space. Each grid cell has peaks in its positional tuning curves at a randomly chosen offset from each lattice point. Its directional tuning curve is uniform. Head direction cells have peaks in their directional tuning curves at a randomly chosen angle and have uniform positional tuning curves. Conjunctive cells have positional tuning curves like grid cells and directional tuning curves like head direction cells. We describe tuning curves for the non-periodic neural populations in the Methods section.

These tuning curves are applied to an experimentally extracted trajectory of a rat exploring its circular enclosure, producing an activity, or firing rate, for each neuron at 0.2 s intervals. This simulates the simultaneous recording of a large number of neurons from the medial entorhinal cortex and the binning of their spikes into firing rates. The time series spans 1000 s, or 5000 data points. Figure 1A shows examples of these time series data mapped back onto spatial coordinates. Next, we choose a subset of these neurons and pre-process it for topological data analysis (Fig. 1B). We form a vector of neural activities at each timepoint, which produces a point cloud in high-dimensional phase space. We wish to subsample it for computation tractability while maintaining as much evidence of topological structure embedded within it. To do so, we use a geometric subsampling algorithm that roughly eliminates the most redundant points, reducing the 5000 timepoints down to 1000.

Finally, we apply persistent cohomology to this subsampled point cloud (Fig. 1C). We describe this technique colloquially here, in terms of its dual, persistent homology. Both produce the same persistence diagrams, but we use cohomology throughout the paper both because it is faster to compute and because it allows us to parametrize the data. See the Methods section for a precise description. From the point cloud, we form a Vietoris–Rips filtration, which is a nested sequence of simplicial complexes. Each complex consists of all cliques in the near-neighbor graph, which contains all edges between points at distance at most *r* apart. As the threshold *r* increases, more edges enter the graph, and more cliques enter the Vietoris–Rips complex. Throughout this process, cycles (e.g., 1-dimensional loops) appear and get filled in the complex (Fig. 2A). There is a unique way to pair the distance thresholds at which cycles are born and die.

**Figure 2:**
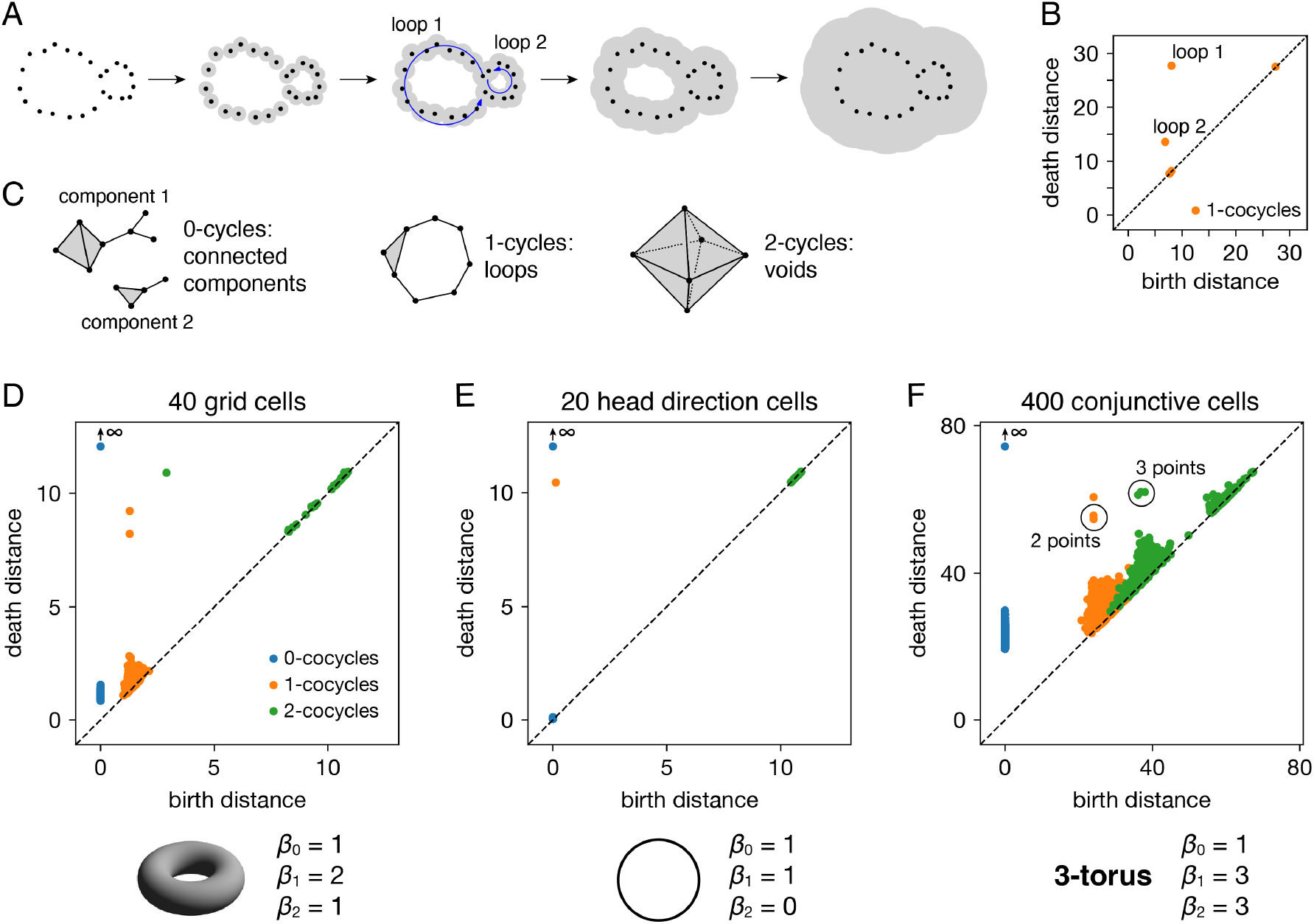
Persistence diagrams for periodic neural populations. (**A–C**) Interpreting persistence diagrams. (**A**) Persistent (co)homology involves generating a filtration of a dataset in which we connect points within ever-expanding distances. One-dimensional loops will be born and then die at various distances. (**B**) These values are plotted in a persistence diagram where each point corresponds to a single loop. The points located farthest from the diagonal are the most persistent and correspond to significant topological features. (**C**) This process can be performed for features of arbitrary dimension *k*. (**D–F**) Persistence diagrams for periodic neural populations (top) and identified topological spaces (bottom). We compare the number of persistent *k*-(co)cycles to the *k*-th Betti numbers *β*_*k*_ of different topological spaces to infer the underlying topological structure of the dataset. (**D**) Grid cells from module 1 exhibit one persistent 0-cocycle, two persistent 1-cocycles, and one persistent 2-cocycle, which corresponds to a torus. (**E**) Head direction cells exhibit one persistent 0-cocycle, one persistent 1-cocycle, and no persistent 2-cocycles, which corresponds to a circle. (**F**) Conjunctive cells exhibit one persistent 0-cocycle, three persistent 1-cocycles, and three persistent 2-cocycles, which corresponds to a 3-torus.

All such birth and death distances are collected into a persistence diagram (Fig. 2B). The points farthest from the diagonal correspond to the most persistent cycles that appear for the longest range of distance thresholds. They recover topological structure in the space sampled by the point cloud, which corresponds to the processes underlying the data—in our case, the spatial representation networks and external inputs. Persistent (co)homology is stable: the persistent points will remain in the diagram if we make small changes to the data, such as selecting slightly different timepoints or perturbing their values by a small amount of noise. The points closest to the diagonal would appear even if the processes underlying the data lack topological structure, and they are usually interpreted as noise.

The process we described above keeps track of cycles of different dimensions (Fig. 2C). Besides loops (1-dimensional cycles), it tracks connected components (0-dimensional cycles), voids (2-dimensional cycles) and higher-dimensional topological features, which lack a colloquial name. The number of independent *k*-cycles is called the *k*-th Betti number and is a topological invariant of a space. We can infer the topology of a dataset by comparing the number of persistent *k*-(co)cycles to the *k*-th Betti numbers of conjectured ideal spaces, such a circle or a torus. Note that for every dataset, the 0-(co)cycle corresponding to the entire point cloud will never die, so we consider its death distance to be infinity.

### 2.2 Persistent cohomology for periodic neural populations

Each periodic neural population spans a particular topological space. We recover these relationships when we compute persistent cohomology of our simulated data (Fig. 2D–F). Each grid cell is active at one location in a unit cell that is tiled over 2D space. Grid cells within a single module share the same unit cell but differ in their active location, so each grid module spans a torus, which is periodic in two directions [5]. Similarly, head direction cells span a circle and each conjunctive cell module spans a 3-torus. The correspondence between our results and predicted topological spaces validates the basic capabilities of our methods.

The ability of persistent cohomology to discover topological structure depends on the number of neurons in the dataset, or equivalently, the dimension of the time series embedding. Using the grid cell population as an exemplar, we form multiple datasets with randomly selected neurons to measure the success rate of persistent cohomology as a function of neuron count (Fig. 3A). To measure success, we only use the first cohomology group H^1^, which contains 1-cocycles. We define successful discovery of the grid cell torus as a persistence diagram with two persistent 1-cocycles, and we define what it means to be persistent precisely using the commonly used *largest-gap heuristic*. We calculate the lifetime of each cocycle, which is the difference between its death and birth and corresponds to the vertical distance to the diagonal of its point in the persistence diagram (Fig. 3A,B). We find the largest gap in the lifetimes and consider points above this gap to be persistent (Fig. 3B). Figure 3C shows that reliable discovery of the torus using this heuristic can be achieved with ≈20 simulated, idealized grid cells. It also shows that increasing the number of grid cells improves the success rate. This occurs because topological discovery relies on having enough grid cells such that their fields provide a sufficiently uncorrelated coverage of the unit cell. Instead of using data spanning the full 1000 s-long trajectory, which corresponds to 5000 timepoints, we extract portions spanning various lengths. Discovery of the torus requires enough samples of its manifold structure, which is achieved in this case starting at 100 s (Fig. 3D). Thus, persistent cohomology generally thrives in the large-dataset limit with long neural recordings of many neurons.

**Figure 3:**
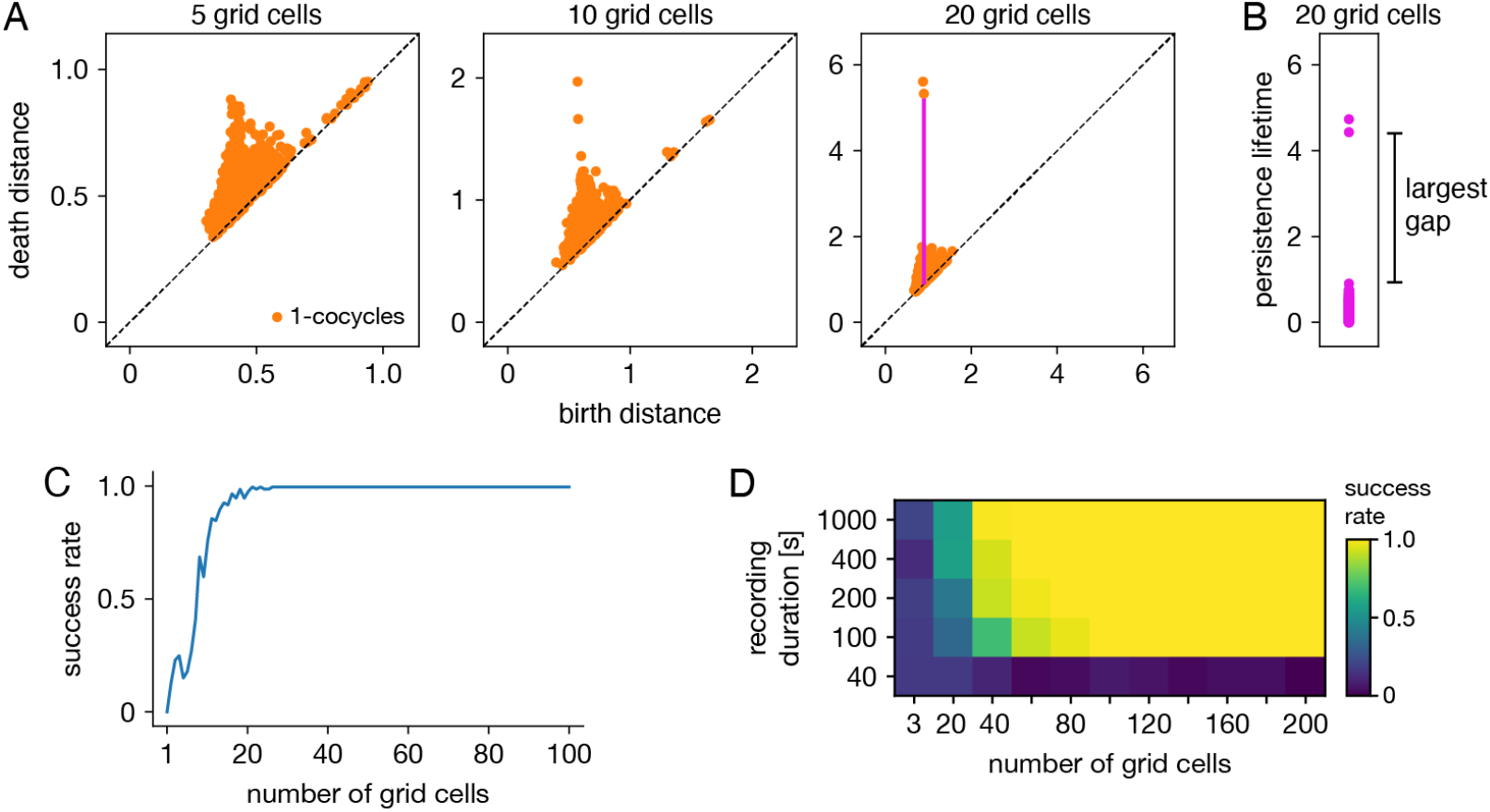
Success rates of persistent cohomology for grid cells. (**A**) As the number of grid cells increases, two persistent 1-cocycles emerge. To define persistence precisely, we consider the persistence lifetime of each 1-cocycle, which is the difference between its birth and death distances (length of the magenta line). (**B**) We identify the largest gap in persistence lifetimes and define persistent cocycles as those above this gap. (**C**) Since a grid module has a toroidal topology for which *β*_1_= 2, we define success as a persistence diagram with two persistent 1-cocycles. We determine success rates by generating 100 replicate datasets. Success rate increases with the number of grid cells. (**D**) Success rates for different durations extracted from the full simulated recording. Topological discovery benefits from longer recording durations and more neurons.

Persistent cohomology can succeed for mixed signals. Separation of raw electrode recordings into single-neuron spike trains may not always be possible or desired. To address this scenario, we form multi-neuron units by linearly combining time series of neural activity across different grid cells. The mixing coefficients are drawn from a uniform random distribution and then normalized. Example activity maps of these multi-neuron units as a function of position are shown in Fig. 4A. The combination of many neurons destroys the classic responses exhibited by individual grid cells. Yet, multi-neuron units retain topological information associated with the grid module that can be recovered by persistent cohomology (Fig. 4B). The success rate for discovering toroidal topology is remarkably independent of the number of grid cells in each unit. Successful recovery from randomly mixed signals is not entirely unexpected. The preservation of distances under random projections has been studied extensively in statistics (cf. Johnson–Lindenstrauss lemma [29]).

Persistent cohomology can also succeed in the presence of spiking noise. To simulate such noise, we use our generated activity as a raw firing rate that drives a Poisson-like random process (see Methods). We construct this process to have different Fano factors, which is the variance in the random process for a given firing rate divided by the firing rate. When the Fano factor is 1, the random process is Poisson. Figure 4C shows activity time series for two grid cells that have very similar tuning curves and thus very similar raw firing rates, which can be seen in the noise-free condition (top left). Higher Fano factors lead to more variability both across time for each neuron and across neurons. Persistent cohomology can still recover the toroidal topology of the grid module, though more neurons are required for higher Fano factors (Fig. 4D). In the mammalian cortex, Fano factors lie around ∼1.0–1.5 [30]. Applying this regime to our simulations implies that ≈80 grid cells are required for reliable topological discovery, but we acknowledge the large differences between simulated and experimental data which may substantially increase this number.

**Figure 4:**
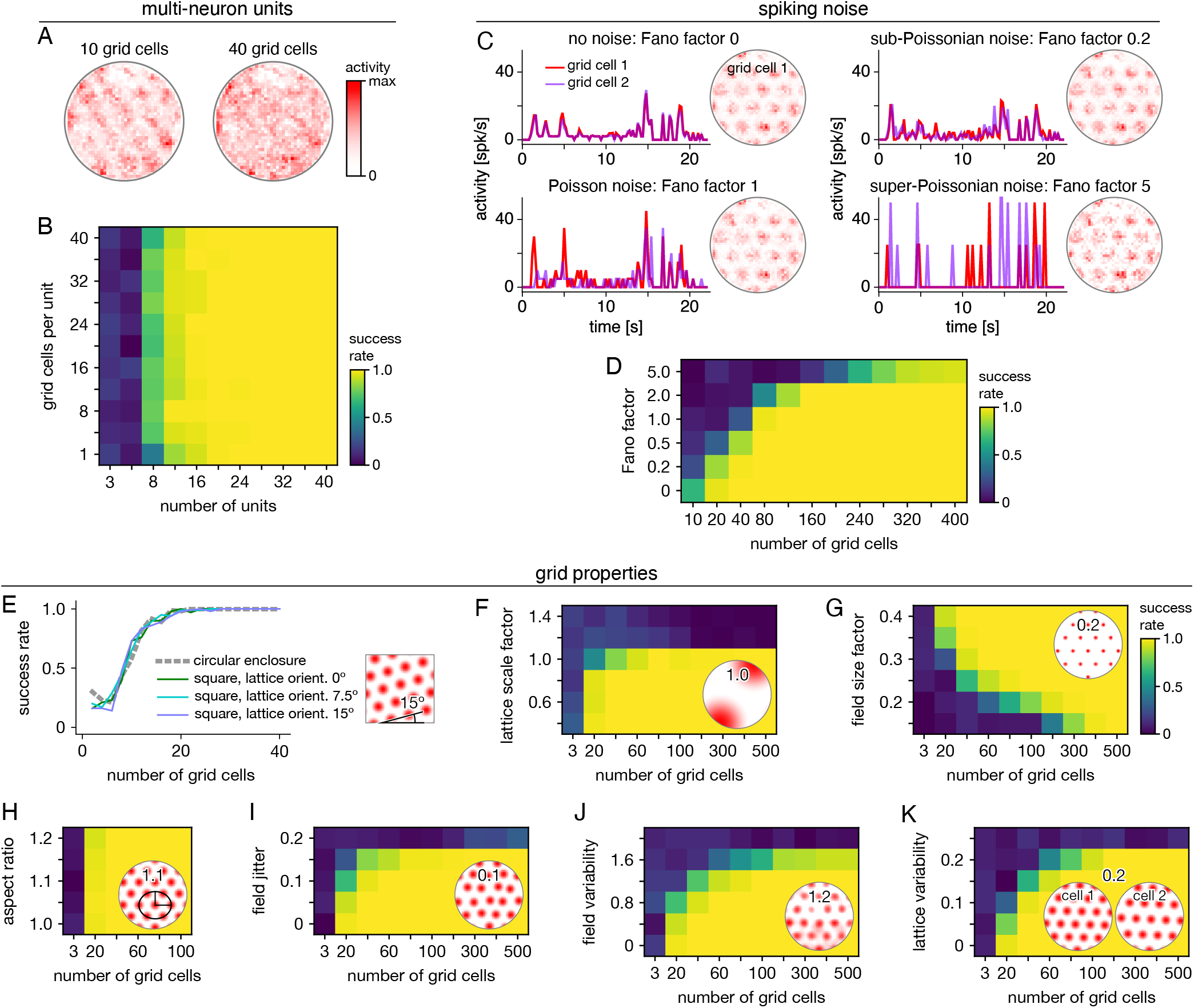
Robustness of persistent cohomology for grid cells. (**A**,**B**) Linearly combining activities from multiple neurons before applying persistent cohomology. (**A**) Example positional activity maps for multi-neuron units. (**B**) The success rate is largely independent of the number of neurons per multi-neuron unit. (**C**,**D**) Introducing Poisson-like spiking noise with different amounts of variability, as measured by the Fano factor. (**c**) For each Fano factor, time series for two grid cells with very similar offsets (left) and positional activity map of the first (right). (**D**) Persistent cohomology requires more neurons to achieve success with higher Fano factors. (**E–K**) Success rates across many different grid cell variables with insets depicting example values. See Methods Sec. 4.1.7 for a complete description of each variable. (**E**) Introducing a square enclosure at various lattice orientations. (**F**) Changing the lattice scale factor, which is grid scale divided by enclosure diameter. (**G**) Changing the field size factor, which is grid field diameter divided by grid scale. (**H**) Stretching triangular grids to various aspect ratios. (**I**) Jittering the position of each field by random vectors multiplied by grid scale. (**J**) Introducing variability in shape and overall activity across grid fields. (**K**) Introducing variability in lattices across grid cells.

We further test the robustness of persistent cohomology across wide ranges of properties associated with grid cells (Figure 4E–K). Using a trajectory extracted from an animal exploring a square enclosure does not meaningfully change the success rate (Fig. 4E). Grid modules have been shown to favor a lattice orientation of 7.5^°^ relative to square enclosures [31], but this orientation does not affect topological discovery. Similarly, persistent cohomology is not strongly affected by the aspect ratio of the triangular grid (Fig. 4H). Changes in grid dimensions can have a stronger effect. Persistent cohomology fails when the grid scale exceeds the size of the enclosure, which makes sense because the grid module unit cell can no longer be fully sampled (Fig. 4F). When the scale does not change but the size of each firing field is decreased, more grid cells are required to produce toroidal structure in the data since each neuron covers less of the unit cell (Fig. 4G). Under various forms of variability within grid cell tuning curves, the success of persistent cohomology can be maintained if more neurons are recorded, up to a degree (Fig. 4I–K). Beyond critical values, however, variability appears to catastrophically disrupt topological discovery in a way that cannot be overcome with more grid cells. The extra heterogeneity conveyed by an additional neuron overwhelms its contribution to topological structure. Thus, an assessment of tuning curve variability may be a crucial component in the application of persistent cohomology to neural data.

### 2.3 Animal trajectory decoded through topological coordinates

Persistent cohomology can not only discover topological structure in neural data, but it can also decode information embedded within this structure. Recall that a grid module defines a triangular lattice in physical space with fields of each grid cell offset in the rhombic unit cell (Fig. 5A). The periodicities of this unit cell along the two lattice vectors are detected by persistent cohomology as two persistent 1-cocycles belonging to a torus (as seen in Fig. 2D). We can assign circular coordinates [15, 32] for these 1-cocycles (Fig. 5B). These coordinates describe the topological space of the torus and should map back onto the rhombic unit cell that tiles physical space. To explore this relationship, we project the entire time series of neural activities onto these coordinates. For each neuron, we find the data points for which that neuron is the most active within the population. These points are clustered and define firing fields in topological space (Fig. 5C). The center of masses of these clusters are used to evaluate distances between grid cells; these topological distances are highly correlated with the physical distances between grid offsets within the rhombic unit cell (Fig. 5D). Thus, the topological coordinates defined in neural activity space indeed capture the organization of grid cells in physical space.

**Figure 5:**
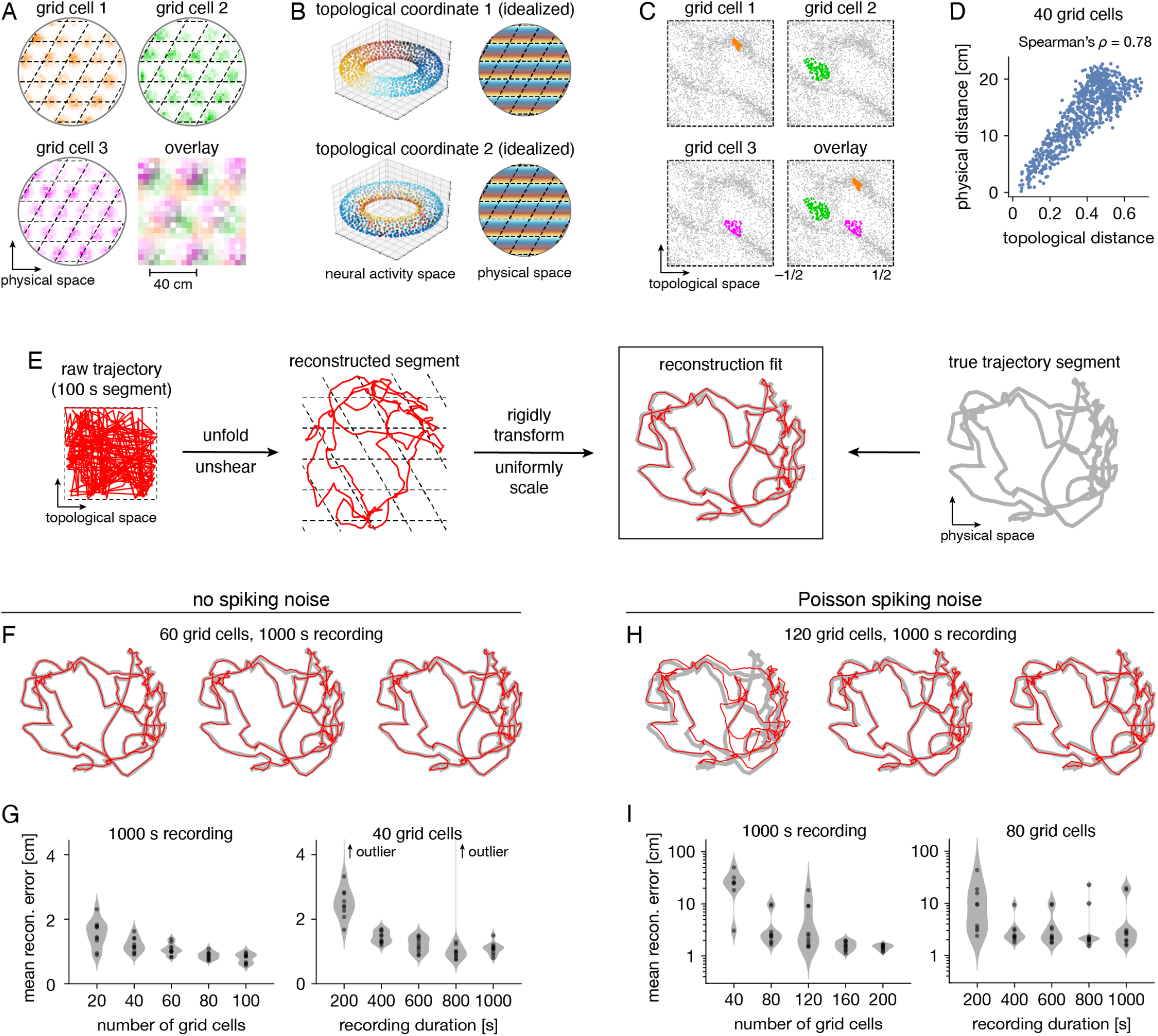
Correspondence between spatial and topological coordinates for grid cells. (**A**) Positional activity maps. A grid module tiles physical space with a rhombic unit cell (black dashed lines) with a different offset for each grid cell. (**B**) Schematic of circular coordinates (color) for the two persistent 1-cocycles of a grid module that define a topological space (left) and their mapping onto physical space (right). (**C**) Firing fields in the topological space of neural activities. All timepoints (gray) and those for which a grid cell is the most active in the population (color). (**D**) Topological distances between firing fields depicted in **C** are correlated with physical distances between grid offsets depicted in **A**. Each point represents distances between two grid cells. (**E**) Reconstructing a segment of animal trajectory in the topological space of neural activities and fitting it to the true trajectory segment. (**F**,**G**) Reconstructions for grid cells without spiking noise. (**F**) Sample reconstruction fits. (**G**) Mean reconstruction error is the distance between reconstructed and actual positions averaged over timepoints. For each condition, we generate 10 replicate datasets, shown as violin plots with points representing individual values. Error generally decreases with increasing grid cell number and trajectory duration. (**H**,**I**) Same as **F**,**G** but for grid cells generated with Poisson spiking noise and error plotted on a logarithmic scale. Geometric subsampling was not performed. Persistent cohomology can achieve low reconstruction error with large enough grid cell number and trajectory duration.

Furthermore, persistent cohomology can leverage the mapping between physical and topological spaces to decode trajectories in the former by trajectories in the latter (Fig. 5E). To do so, we trace the circular coordinates depicted in Fig. 5C through time to form a raw trajectory through topological space. We then unshear it by 60° and unfold this trajectory by identifying large jumps with wrapping through the boundary to produce a reconstructed segment. These steps do not require knowledge of the animal’s true trajectory (although some general expectations about the animal’s motion are required for unshearing; see Methods Sec. 4.5 for details). We find that this reconstruction can be translated, rotated, reflected, and/or uniformly scaled to match the true trajectory very well. Without spiking noise, almost all reconstructions deviate from the true trajectory by less than 4 cm averaged across time, which is much less than the enclosure’s diameter of 180 cm (Fig. 5F,G). This error decreases with more neurons or more timepoints in the simulated recording. Similar trends are observed with the introduction of Poisson noise that mimics spiking noise, but more neurons are required, more outliers with poor fit are observed, and geometric subsampling cannot be used to improve computational tractability (Fig. 5H,I).

### 2.4 Persistent cohomology for mixtures of neural populations

Persistent cohomology can discover topological structure in mixtures of neural populations. When neurons are recorded from a periodic neural population and a non-periodic neural population, the latter adds additional dimensions to the point cloud embedding, but the topological structure contained within the former should persist. We test if persistent cohomology can recover this information in mixed datasets with neurons from both a periodic population (either grid or conjunctive) and a non-periodic population (either non-grid spatial or random). Reliable discovery of the torus formed by grid cells is possible when the number of spatial or random cells is less than twice the number of grid cells (Fig. 6A,B). Detection of the 3-torus formed by conjunctive cells requires more neurons, but it can also be reliably achieved in the presence of non-periodic populations (Fig. 6C,D). Thus, persistent cohomology demonstrates robustness to the inclusion of non-periodic populations. The size of the non-periodic population that can be tolerated increases with the size of the periodic population.

**Figure 6:**
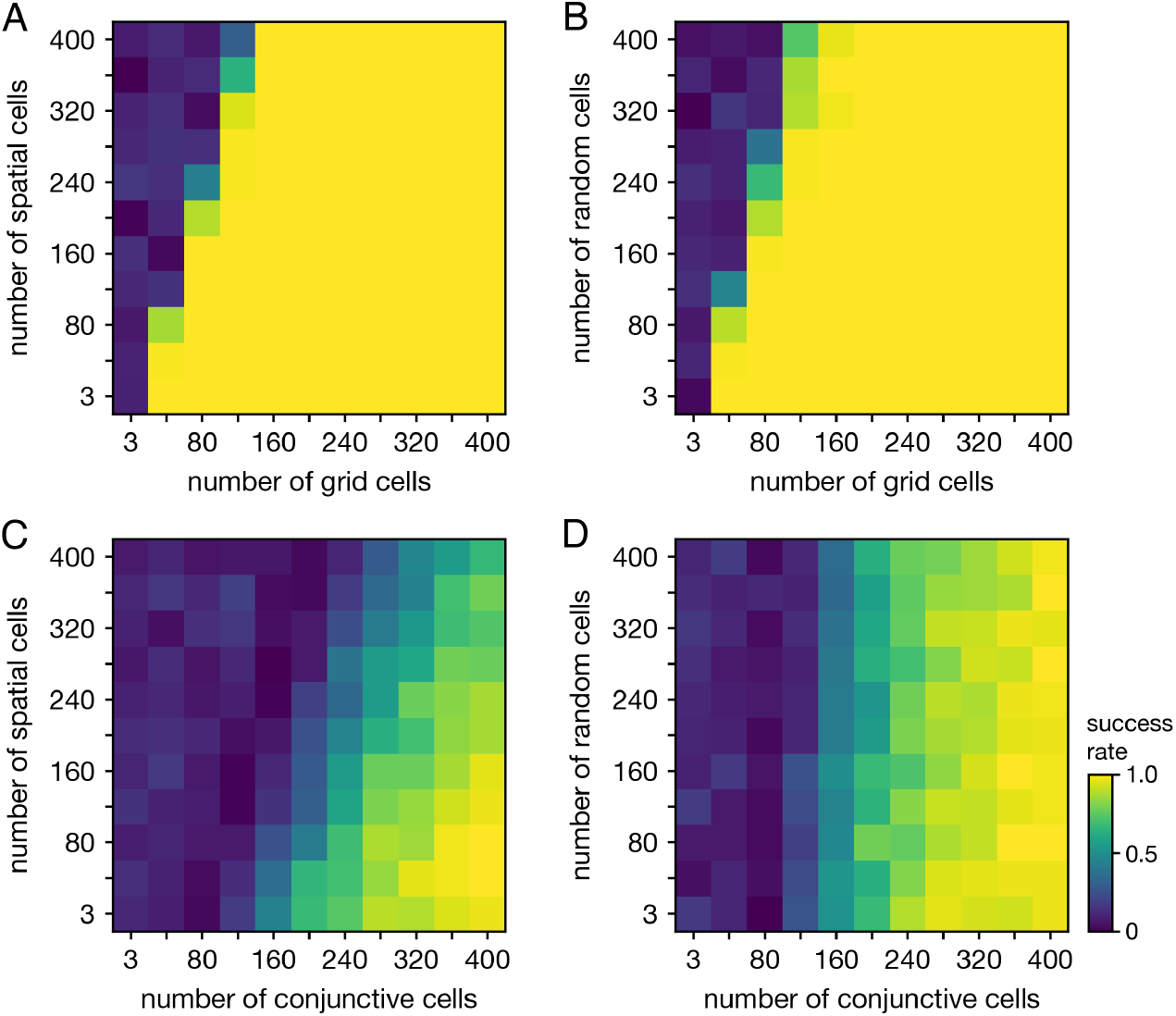
Persistent cohomology in combinations of periodic and non-periodic neural populations. Success is defined by observing the number of persistent 1-cocycles expected from the periodic population, which is two for grid cells and three for conjunctive cells. (**A**) Grid cells from module 1 and non-grid spatial cells. (**B**) Grid cells from module 1 and random cells. (**C**) Conjunctive cells from module 1 and non-grid spatial cells. (**D**) Conjunctive cells from module 1 and random cells.

When neurons are recorded from multiple periodic neural populations, their structures are preserved along separate subspaces in high-dimensional activity space. We explore this scenario by forming mixed datasets with neurons from two periodic populations. When the two populations respond to unrelated signals—such as grid and head direction cells—the combined topological space should be the Cartesian product of those of the separate populations. Indeed, that persistent cohomology can discover the resultant 3-torus at intermediate mixing ratios (Fig. 7A,B). If one population contributes many more neurons—and thus embedding dimensions—than the other, we instead detect the corresponding single-population structure (Fig. 7B).

**Figure 7:**
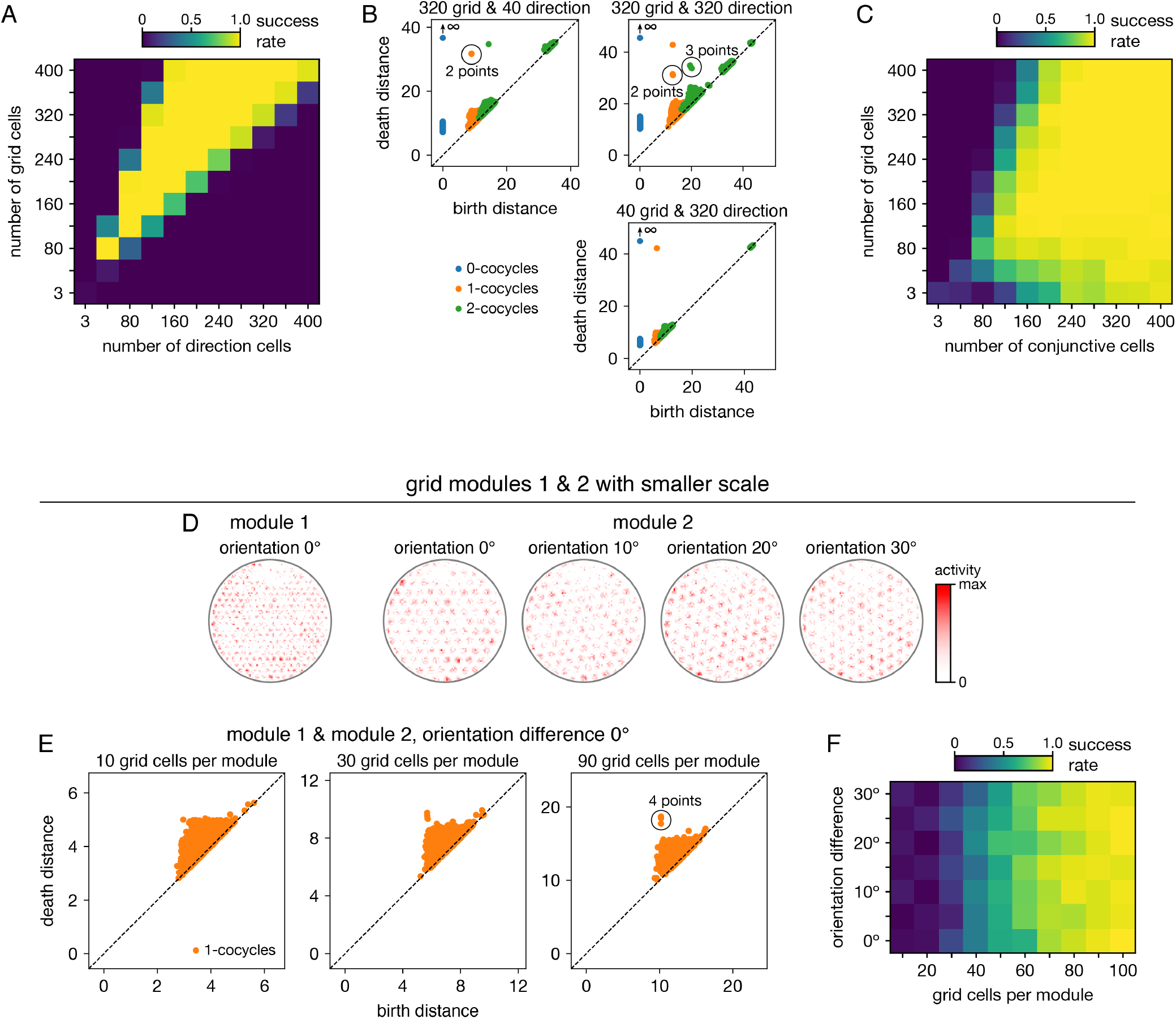
Persistent cohomology in combinations of periodic neural populations. Success is defined by observing the number of persistent 1-cocycles expected from the product topology. (**A**,**B**) Discovering the 3-torus for grid cells from module 1 and head direction cells. (**A**) Success rates. (**B**) Example persistence diagrams. If one population contributes many more neurons than the other, persistent cohomology detects the topology of the predominant population. (**C**) Success rates for discovering the 3-torus for grid cells and conjunctive cells, both from the same module 1. (**D–F**) Combining grid cells from two modules with smaller scales. (**D**) Positional activity maps for example neurons from module 1 and four different orientations of module 2. (**E**) Example persistence diagrams. (**F**) Success rates for discovering the 4-torus.

When the two populations respond to related signals—such as grid and conjunctive cells—the activity space of one is contained in the activity space of the other. Grid cells and conjunctive cells from the same module encode position with the same toroidal structure; they both tile space with the same rhombic unit cell of neural activity. In addition, the conjunctive population encodes direction with a circular topology. Thus, the mixed dataset should span a 3-torus, which can be detected by persistent cohomology (Fig. 7C). For reliable discovery, at least ≈120 conjunctive cells and at least ≈240 total neurons are required. However, discovery of the product topology is disrupted if the number of grid cells exceeds the number of conjunctive cells by more than a factor of ≈1.5. Thus, persistent cohomology can best detect product topologies when the mixed dataset is not dominated by one population.

Finally, we consider the case of mixing grid cells from multiple modules. Grid modules have different rhombic unit cells with different scales and orientations, so they map the same physical space onto different topological coordinates. Thus, a mixed dataset from two different modules should exhibit the product topology of two 2-tori, which is the 4-torus. However, we are unable to reliably discover this structure using the grid modules illustrated in Fig. 1A; they are too sparse. To produce a point cloud that embeds the toroidal structure for one grid module, the animal trajectory should densely sample its rhombic unit cell. This is achieved since the enclosure contains many unit cells. However, to produce a point cloud that embeds the 4-torus formed by two grid modules, the animal trajectory should densely sample all combinations of unit cells. This is not achieved by the grid modules illustrated in Fig. 1A because the enclosure contains too few rhombic unit cells for them to overlap in many different configurations.

Thus, we generate two grid modules separated by the same scale ratio as in Fig. 1A, but with one-fourth of its scale (Fig. 7D). In addition, we explore different relative orientations between the modules by generating different orientations for module 2. Notably, these scale ratios and orientation differences are not chosen such that the two rhombic unit cells would share a simple geometric relationship with each other [33], which would limit their possible overlap configurations and collapse the expected 4-torus structure to a 2-torus. As we include more neurons from both modules into our dataset, we see that four persistent 1-cocycles eventually emerge from the points close to the diagonal that represent sampling noise (Fig. 7E). The success of persistent cohomology is independent of the orientation difference between the two modules (Fig. 7F). Note that if we obtained independent activity samples from each module, combined datasets formed from the original grid modules with larger scales should exhibit 4-torus structure. However, samples taken from a single animal trajectory are not independent across modules, so smaller grid scales (or equivalently, larger environments) are required to fully sample the 4-torus structure.

## 3 Discussion

We demonstrate that persistent cohomology can discover topological structure in simulated neural recordings with as few as tens of neurons from a periodic neural population (Fig. 3). From this structure, it can decode the trajectory of the animal using only the time series of neural activities (Fig. 5). It can also discover more complex topological structures formed by combinations of periodic neural populations if each population is well-represented within the dataset (Fig. 7).

By comprehensively adjusting a wide range of parameters related to grid cells, we find that persistent cohomology generally behaves in three different ways with respect to parameter variation. First, topological discovery can be unaffected by some manipulations, such as combining grid cells into multi-neuron units and changing global geometric features such as enclosure geometry and lattice aspect ratio (Fig. 4). Second, topological discovery may be impeded in a way that is counteracted by increasing neuron number. Spiking noise, small field sizes, and inclusion of non-periodic populations are examples of parameters that exhibit this behavior (Figs. 4 and 6). Third, topological discovery can fail catastrophically in certain parameter regimes without the possibility of recovery by including more neurons. This happens if tuning curves are inherently too variable or if discovery of product topologies are desired when one neural population vastly outnumbers the other (Figs. 4 and 7). These conclusions can help researchers understand and overcome obstacles to topological discovery with persistent (co)homology and may guide its use across a variety of neural systems.

We have characterized the capabilities of persistent cohomology using simple simulated data, but our results may generalize to real neural data. The inputs to our analysis pipeline are firing rates over 0.2 s time bins, which averages over many neurophysiological processes, including major neural oscillations found in the hippocampal region [34]. Moreover, we have found that persistent cohomology is robust to many simulated sources of biological noise. A key requirement for generalization is a separation of two timescales. The macroscopic timescale at which topological structures are explored—here, the time required to traverse a rhombic unit cell of a grid module or 360^°^ of head direction—must be much longer than the microscopic timescale at which neuronal activity is generated. This enables us to coarse-grain over the activity and describe it by a firing rate.

For comparison, we attempt to modify manifold learning algorithms to enable discovery and interpretation of topological structure (Supplementary Material and Supp. Fig. 1). We find that Isomap [35] followed by Independent Components Analysis (ICA) [36] can successfully identify and decode from toroidal structure with modified grid cells whose tuning curves exhibit a square lattice. Unlike persistent cohomology, it does not work for triangular lattices and does not consider topological features with dimensionality greater than 1. UMAP [37] can be used to embed grid cell data directly into a 2-dimensional space with a toroidal metric, but we find that its coordinates do not correspond well to the physical grid periodicity. In short, persistent cohomology performs better and requires adjusting fewer parameters for topological data analysis compared to these alternative methods, which were not designed for such analysis. Note that [25] formulate an alternative method for decoding topologically encoded information; they also use persistent homology for discovery of topological structure.

The application of persistent (co)homology to neuroscience data is still in its developing stages. In addition to the research on spatial representation circuits described in the Introduction [24, 25, 26, 27], notable lines of work include: simulations of hippocampal place cells in spatial environments with nontrivial topology [38, 39, 40, 41, 42]; analysis of EEG signals, for classification and detection of epileptic seizures [43, 44] and for construction of functional networks in a mouse model of depression [45]; inferring intrinsic geometric structure in neural activity [46]; and detection of coordinated behavior between human agents [47]. There is potential for persistent (co)homology to provide insight to a wide range of neural systems. Topological structures generally can be found wherever periodicities exist. These periodicities can take many forms, such as the spatial periodicities in our work, temporal regularities in neural oscillations, motor patterns, and neural responses to periodic stimuli.

The toolbox of topological data analysis has more methods beneficial to the analysis of neural data. The methods described in this paper, including geometric subsampling, are sensitive to outliers. This problem can be addressed within the same framework of persistent cohomology by using the distance-to-a-measure function [48]. In practice, this would translate into a slightly more elaborate construction [49] of the Vietoris– Rips complex. Furthermore, our analysis pipeline benefits from having neural activity embedded in a high dimensional space, i.e., from having many more neurons than the intrinsic dimensions of the recovered tori. It is possible to adapt this technique to the regime of limited neural recordings (even to a single neuron) by using time-delay embeddings [50]. However, for spatial populations, such a technique would require control over the smoothness of the animal’s trajectory, which may not be feasible in practice. Also, the method we present cannot make inferences on network topology. If connectivity information were present in neural activities, they should appear on fast timescales related to synaptic integration, action potential propagation, and synaptic delay. By averaging neural activity into 0.2 s time bins, we destroy this information, but it is possible that a modified method may access it.

Our results also suggest research directions in topological data analysis. Throughout the paper, we relied on 1-dimensional persistent cohomology to infer whether we recovered a particular torus. But that is a relatively weak method: many topological spaces have cohomology groups of the same dimension. Although the trajectories that we recover via circular coordinates serve as a convincing evidence that we are indeed recovering the tori, it is possible to confirm this further by exploiting cup product structure in cohomology, which is a particular kind of a topological operation that turns cohomology into a ring. Computing a “persistent cup product” would provide additional evidence about the structure of the recovered spaces.

## 4 Methods

### 4.1 Generating neural recordings

#### 4.1.1 Animal trajectory

We simulate the simultaneous recording of neurons from a rat as it explores a circular enclosure of diameter 1.8 m. We use 1000 s from a trajectory extracted from an experimental animal [16, 51]. This trajectory is provided as velocities sampled at 0.5 ms intervals along with the initial position. We average these to positions and directions at 0.2 s intervals as follows: the position of the animal is simply the average position within each 0.2 s time bin, and the direction of the animal is the circular mean of the velocity vector angle within each 0.2 s time bin. We ignore the distinction between body direction and head direction.

#### 4.1.2 Periodic neural populations

We generate tuning curves as a function of position and/or direction for each neuron. These localized tuning curves are based on a shifted and truncated cosine function:

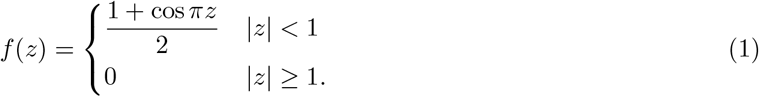

For each grid module, we set a scale *l* and an orientation *ϕ*. This defines a transformation matrix from the space of phases 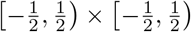 to the rhombic unit cell of the grid module in physical space:

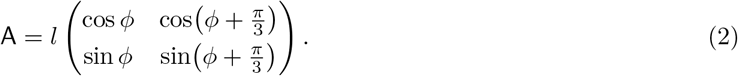

Unless otherwise specified, we use *l* = 40 cm and *ϕ* = 0. The inverse of this matrix A^−1^ maps the rhombic unit cell onto the space of phases. We also define ‖ · ‖ as the vector norm and ⟨*a*⟩_*m*_ ≡ (*a* + *m* mod 2*m*) − *m* as the shifted modulo operation. The tuning curve of a grid cell as a function of position **x** is then

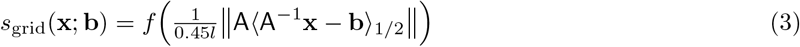

Each grid cell is shifted by a uniformly random phase offset **b**. The full width at half maximum of each grid field is 0.45*l*. In physical space, the offset of a grid cell is A**b**, where integers can be added to either component of **b**; the shortest distance between these offsets for two grid cells is the physical distance shown in Fig. 5D.

The tuning curve of a head direction cell as a function of direction *θ* is

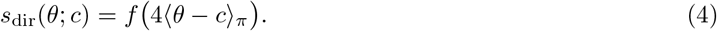

Each head direction cell is shifted by a uniformly random direction offset *c*. The full width at half maximum of the head direction field is *π/*2.

The tuning curve of a conjunctive cell is simply the product

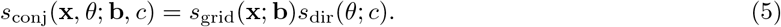

Each conjunctive cell has uniformly random offsets **b** and *c*.

#### 4.1.3 Non-periodic neural populations

For non-grid spatial cells, we generate tuning curves as a function of position for each neuron of the form

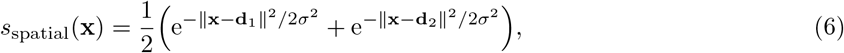

where σ = 40 cm and the **d**_*i*_’s are chosen uniformly randomly between (0 cm, 0 cm) and (180 cm, 180 cm).

For random neurons, we obtain activity time series by sampling from a distribution every 2 s, or 10 timepoints, and interpolating in between using cubic polynomials. The distribution is Gaussian with mean 0 and width 0.5, truncated between 0 and 1.

#### 4.1.4 From tuning curves to time series

To obtain activity time series for all populations except for random neurons, we apply the tuning curves to each trajectory timepoint. Whenever the velocity decreases below 5 cm*/*s, we set the activity to be 0. This threshold simulates the behavior of neurons in the hippocampal region that exhibit high activity during locomotion and low activity during idle periods [20, 52, 53].

#### 4.1.5 Multi-neuron units for grid cells (Fig. 4A,B)

We generate multi-neuron units (Fig. 4) by linearly combining activity time series from multiple grid cells. Each mixing coefficient is chosen from a uniform random distribution between 0 and 1. The activity is then normalized by the sum of squares of the mixing coefficients.

#### 4.1.6 Spiking noise for grid cells (Fig. 4C,D)

The activities described above are dimensionless, and we typically do not need to assign a scale because we divide each time series by its mean before applying persistent cohomology. To create spiking noise, however, we must set the firing rate. We linearly rescale the rate given by Eq. 3:

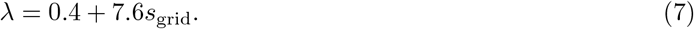

This sets the maximum firing rate to be 8 and creates a baseline rate of 0.4; with 0.2 s time bins, these values correspond to 40 Hz and 2 Hz, respectively. However, we still set the firing rate to 0 Hz when the animal’s velocity decreases below 5 cm*/*s.

Using *λ*, we generate Poisson-like spiking noise with different levels of variability (Fig. 4). At each timestep, the noisy activity is given by

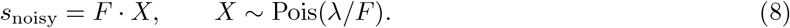

*F* sets the Fano factor of the random process, which is its variance divided by its mean (for any given *λ*). The *F* = 1 case corresponds to a Poisson random process; *F <* 1 implies sub-Poissonian noise and *F >* 1 implies super-Poissonian noise.

#### 4.1.7 Generating grid cells with various properties (Fig. 4E–K)

- Square enclosure (Fig. 4E): We use 1000 s from a trajectory extracted from an experimental animal in a square enclosure of width 1.5 m [17], binned in the same way as for the circular enclosure (Sec. 4.1.1). To change the lattice orientation, we use a nonzero value for *ϕ* in Eq. 2.
- Scale factor (Fig. 4F): Grid scale is modified by changing *l* in Eq. 2. The scale factor is *l* divided by the diameter of the enclosure, 1.8 m.
- Field size factor (Fig. 4G): Field size is modified by replacing 0.45 in Eq. 3 by the field size factor. In other words, the field size factor multiplied by the grid scale is the full width at half maximum of each grid field.
- Aspect ratio (Fig. 4H): The grid lattice is elongated to an aspect ratio *ε* ≥ 1, which is the ratio between the major and minor axes of the ellipse that circumscribes each hexagonal lattice domain. This is accomplished by replacing the transformation matrix in Eq. 2 with

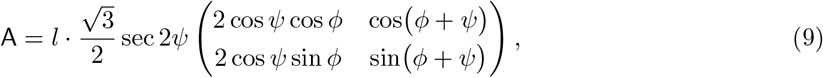

where *ψ ≤* 60^°^ is the angle between the two lattice vectors. *ψ* is related to *ε* by

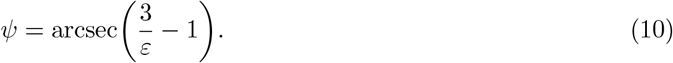

When 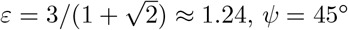 and the lattice becomes square.
- Field jitter (Fig. 4I): Jitter in the position of grid fields is introduced by shifting each field in physical space by

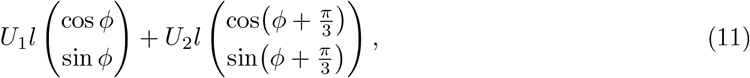

where *l* is the grid scale, *ϕ* is the grid orientation, and *U*_*i*_ ∼ Unif(−field jitter, field jitter).
- Field variability (Fig. 4J): Variability across grid fields is introduced by choosing random numbers *U*_*ij*_ ∼ Unif(−field var., field var.) at 20 cm-intervals throughout the enclosure for each grid cell. A linear interpolation is constructed for these points, which is smoothed by a Gaussian filter of width 20 cm. This resulting random function is added to the standard tuning curve in Eq. 3, and all values under 0 are clipped to 0.
- Lattice variability (Fig. 4K): Variability in lattices across grid cells is introduced by randomly perturbing the transformation matrix for each grid cell (Eq. 2) according to

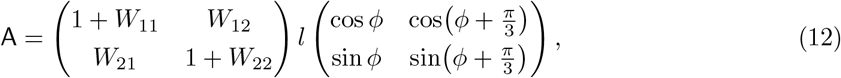

Where

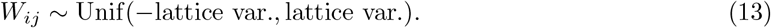

### 4.2 Neural activity maps

We construct activity maps for each neuron as a function of position or direction. To do so, we simply tally the total amount of activity in each positional or directional bin. Note that these maps do not depict firing rate because we do not divide by the occupancy of each bin; we decided against this in order to show the activity experienced through the animal trajectory.

### 4.3 Processing neural recordings

For each neuron, we first divide its activity at every timepoint by its mean activity. We then delete all timepoints whose neural activities are all smaller than a small limit 1 × 10^−4^. This simple procedure removes points at the origin that we do not expect to participate in topological structures.

To improve computational efficiency, we reduce the number of input points while preserving their topological structure by applying a geometric subsampling strategy. We pick the first point at random, and then iteratively add a point to our subsample that is the furthest away from the already chosen points. Specifically, if *P* is the input point set and *Q*_*i*_ is the subsample after *i* iterations, we form *Q*_*i*+1_ by adding the point *q*_*i*+1_ chosen as

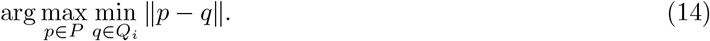

Fig. 1B illustrates a result of this strategy. By construction, the subsample *Q*_*i*_ forms an *ε*_*i*+1_-net of the input point sample, which means the largest distance from any input point to the nearest point of the subsample does not exceed 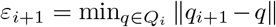. Because persistent cohomology is stable, this guarantees that the persistence diagram we compute for the subsample *Q*_*i*_ is at most *ε*_*i*+1_ away, in bottleneck distance [54], from the persistence diagram of the full point set *P*. We generally select 1000 points through this process. The results in Fig. 5H,I were obtained by applying persistent cohomology on the full dataset without subsampling.

To measure success rate, we apply persistent cohomology on 100 replicate datasets and measure the proportion of successes as determined by the largest-gap heuristic.

### 4.4 Applying persistent cohomology

We refer the reader to extensive literature on persistent (co)homology [13, 14] for the full details. More details on the involved constructions are presented in the Supplementary Material. Here, we only briefly mention some of them. For technical reasons—both to recover the circular coordinates and for computational speed—we work with persistent cohomology, which is dual to persistent homology, which a reader might be more familiar with.

To recover the topology of the space sampled by a point set *P*, we construct a Vietoris–Rips simplicial complex. Given a parameter *r*, Vietoris–Rips complex consists of all subsets of the point set *P*, in which every pair of points is at most *r* away from each other,

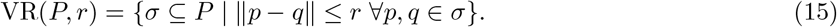

The cohomology group, H^*k*^(VR(*P, r*)), defined formally in the Supplementary Material, is an algebraic invariant that describes the topology of the Vietoris–Rips complex. Its rank, called the *k*-th Betti number, counts the number of “holes” in the complex.

As we vary the radius *r* in the definition of the Vietoris–Rips complex, the simplicial complexes nest: VR(*P, r*_1_) ⊆ VR(*P, r*_2_), for *r*_1_ *≤ r*_2_. The restriction of the larger complex to the smaller induces a linear map on cohomology groups, and all such maps form a sequence:

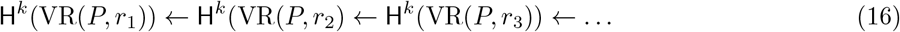

It is possible to track when cohomology classes (i.e., “holes”) appear and disappear in this sequence. Recording all such birth–death pairs (*r*_*i*_, *r*_*j*_), we get a persistence diagram, which completely describes the changes in the sequence of cohomology groups.

For a persistent class, i.e., one with a large difference between birth and death, [15] describe a procedure for turning it into a map from the input data points into a circle. [32] extends that procedure to allow computation of persistent cohomology on a subsample of the data, e.g., the geometric subsample mentioned in the previous subsection.

### 4.5 Reconstructing animal trajectory from circular coordinates

No matter the recording duration used to obtain circular coordinates, we only reconstruct the first 100 s of the animal trajectory. At each timepoint *t* = 1, …, *T*, our circular coordinates form a vector **u**_*t*_ = (*u*_*t*1_, *u*_*t*2_) with components between 0 to 1.

We first perform a preliminary unfolding of the circular coordinates. We calculate all the difference vectors between adjacent timepoints and cancel out changes by more than *±*1*/*2:

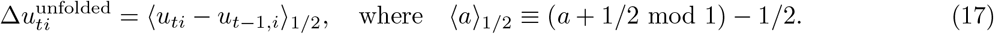

Next, we seek to unshear the coordinates. The rhombic unit cell in physical space is sheared by 30^°^ relative to the orthogonal axes of topological space. We wish to apply this transformation to the difference vectors to restore the unsheared trajectory. There are two possible unshearing matrices

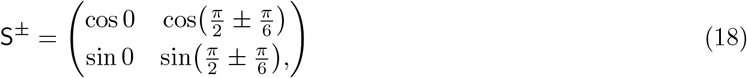

and we could perform the rest of the analysis for both transformations separately, knowing that one trajectory is unsheared and the other is doubly shared. Instead, we assume knowledge that the animal is exploring an open field environment in which all directions of motion should generally be sampled uniformly. We calculate the covariance matrix for both sets of transformed difference vectors:

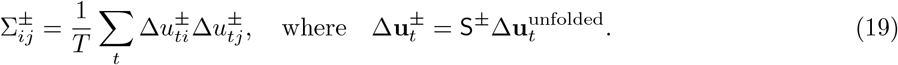

The proper unshearing S produces the covariance matrix whose ratio of eigenvalues is closest to 1. This heuristic could be changed assuming different statistics of animal motion, for example those corresponding to a linear track.

After identifying the unshearing matrix S, we return to the raw coordinates and apply this transformation first, before unfolding:

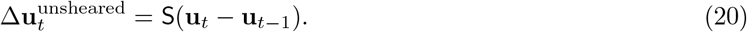

Since shear transformations change distances, we reevaluate our unfolding. We compare the norm of every difference vector 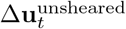 to its norm after possible unfoldings along the unsheared lattice vectors given by the columns of S. The shortest vector at each timepoint is the reconstructed difference 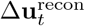, and the reconstructed trajectory segment shown in Fig. 5E is their accumulation 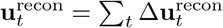

So far, this reconstruction has not incorporated detailed information about the animal trajectory. To judge its quality, we now fit the reconstruction to the true segment of animal trajectory **x**_*t*_ through rigid transformation and uniform scaling. We first determine whether or not the reconstruction needs to be unreflected. To judge this, we calculate the signed vector angles between adjacent steps for both the reconstruction and the true trajectory

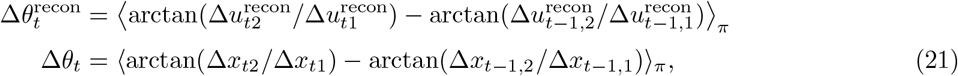

where ⟨*a*⟩_*π*_ ≡ (*a* + *π* mod 2*π*) − *π*. If the mean square difference between 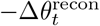 and Δ*θ*_*t*_ is less than that between 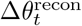 and Δ*θ*_*t*_, then we reflect our reconstruction 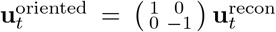. Otherwise,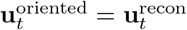.

Finally, we fit the reconstruction to the true trajectory segment by minimizing the mean squared error after uniform scaling *a*, translation **b** and rotation *θ*:

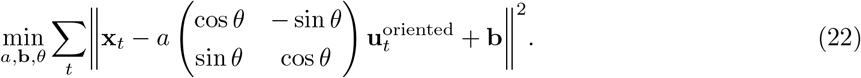

## Acknowledgments

We would like to thank Francesco Fumarola for his insightful and helpful suggestions. L.K. has been supported by a Miller Research Fellowship from the Miller Institute for Basic Research in Science at the University of California, Berkeley. B.X. is supported by a U.S. Department of Energy Computational Science Graduate Fellowship. This work was partially supported by the U.S. Department of Energy under Contract Number DE-AC02-05CH11231 at Lawrence Berkeley National Laboratory.

