## Supplementary Info. and Fig. 1 for "Evaluating state space discovery by persistent cohomology in the spatial representation system"

February 9, 2021

*
†
‡

### 1 Comparison of persistent cohomology to manifold learning algorithms

#### 1.1 Isomap results

To test an alternative method for discovering topological structure, we explored the Isomap manifold learning algorithm [1]. It performs dimensionality reduction by accounting for the geodesic distance along the manifold embedded in high-dimensional space. We tested this method on datasets with a grid module with 40 simulated cells, which exhibits toroidal structure. To visualize the data, we first chose the target dimension to be 3 and observed loops in the point cloud (Supp. Fig. 1A).

To extract the topological structure more rigorously, we performed Isomap with target dimension 4, which is the lowest dimension that permits flat embeddings of a torus. Since Isomap does not output topological information explicitly, we performed Independent Components Analysis (ICA) decomposition on the Isomap embeddings using the FastICA algorithm [2]. ICA identifies the most non-Gaussian directions in the data, which we hoped would participate in 1-dimensional loops in the data, if they exist.

Application of this procedure to simulated grid cells with classic triangular grids revealed a loop embedded in one pair of ICA components but not the other (Supp. Fig. 1B, right). Thus, this dataset would be erroneously identified as possessing circular topology, not toroidal. We found that the ability to detect a second loop in the other pair of ICA components varied with the lattice angle of the simulated grid cells. For square lattices with lattice angle 90°, loops in each pair of ICA coordinates could easily be seen (Supp. Fig. 1B, left). As the lattice angle decreases to the triangular lattice value of 60°, the circular distributions of datapoints become annular and eventually close up.

For ICA decompositions with two loops, we can define circular coordinates for each loop (Supp. Fig. 1B) and use these to reconstruct a segment of animal trajectory, as in Fig. 5 of the main text. We find that Isomap followed by ICA produces accurate reconstructions with errors similar to those obtained by persistent cohomology for square lattices (Supp. Fig. 1C). For triangular lattices, most attempts produced zero or one loop in the data, but occasionally, two loops were revealed (Supp. Fig. 1D). Circular coordinates assigned to these loops could capture the true trajectory of the simulated animal with slightly greater error.

It is possible that toroidal structure is robustly present in the Isomap embeddings of triangular lattice data and that our ICA decomposition cannot reliably detect it. We have explored different ICA contrast functions, as well as Independent Subspace Analysis (ISA) with subspace dimension 2 [3], but could not improve our results. In summary, Isomap can be used to detect and interpret topological structure for some sets of simulated grid cell data. But because it was not designed to perform topological data analysis, it requires more tuning and post-processing than persistent cohomology does.

#### 1.2 UMAP results

We also test the UMAP manifold learning algorithm [4] as an alternative method for interpreting topological structure. We also tested this method on datasets with a grid module with 40 simulated cells, which exhibits toroidal structure. UMAP is often used for visualization, and we find that loops in the point cloud can be observed with target dimension 3 (Supp. Fig. 1E; previously observed by Ref. 5). We do not further assess its ability to detect topological structure; instead, we assume knowledge of the toroidal structure

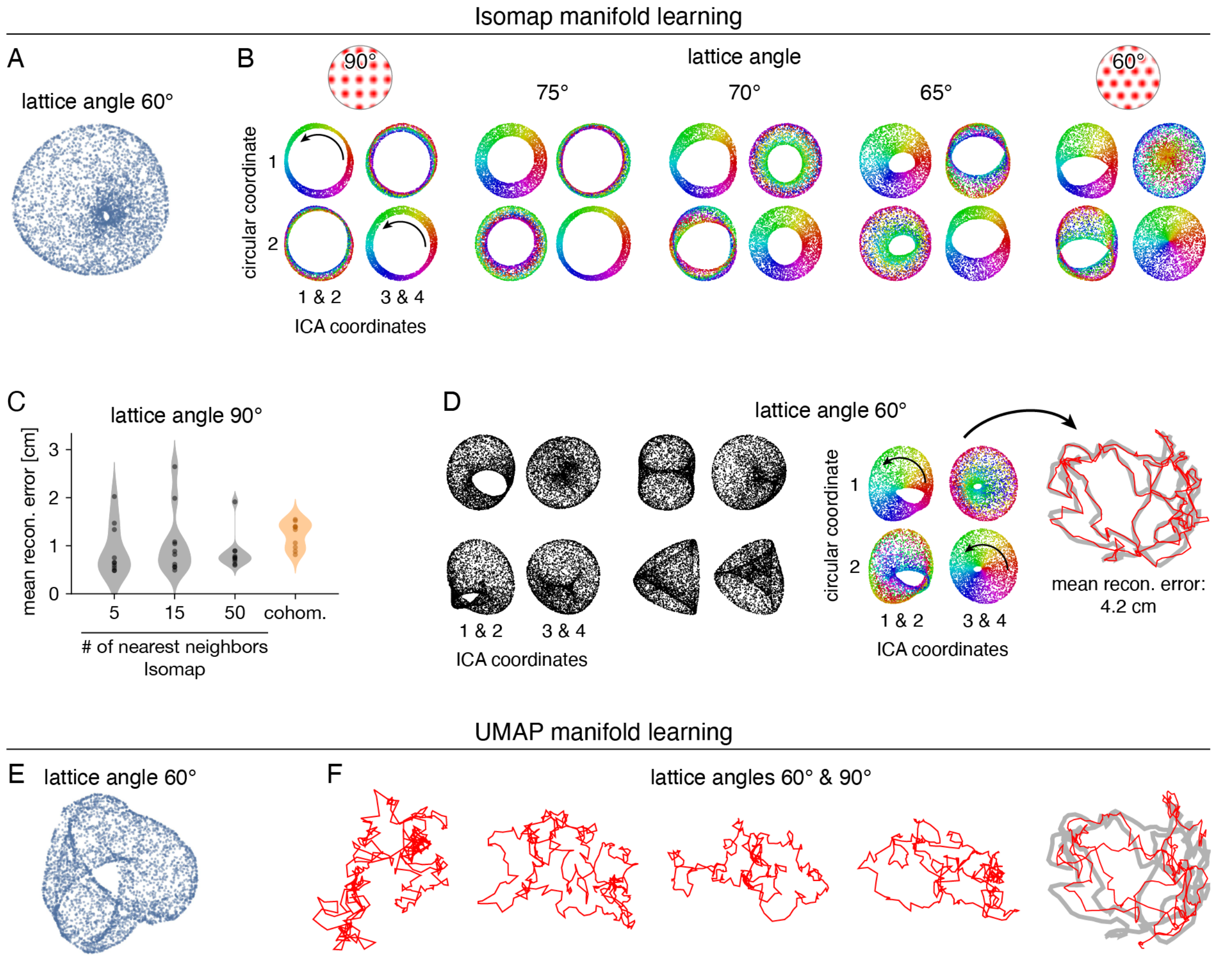

**Supplementary Figure 1**: Topological discovery and circular coordinates with Isomap and UMAP. Analyses performed for datasets with 40 simulated grid cells. We explore not only triangular lattices with lattice angle 60°, but also square lattices with angle 90° and rhombic lattices with other angles. (**A**) Isomap embedding of timepoints into target dimension 3 for visualization. (**B**–**D**) Isomap embedding into target dimension 4, followed by ICA decomposition. (**B**) For grid-like cells with square lattices, the data can be projected onto two sets of two ICA components, each of which exhibits a circular distribution for which a circular coordinate can be defined (rainbow color). As the lattice angle decreases, the circularity decreases. (**C**) The circular coordinates defined for square lattices can be used to reconstruct the animal trajectory with error comparable to that obtained through persistent cohomology. Distributions over 10 replicate datasets are shown as violin plots with points indicating individual values. (**D**) Usually for triangular lattices, circular coordinates cannot be recovered due to noncircular ICA projections. Occasionally, both projections contain annular distributions from which we can define circular coordinates and perform trajectory reconstruction. Reconstructed trajectory segments (red) are fit to true segments (gray). (**E**) UMAP embedding of timepoints into target dimension 3 for visualization. (**F**) Reconstructed trajectories for UMAP embedding of timepoints onto target dimension 2 using a toroidal distance metric. Reconstructed trajectory segments (red) seldom fit true segments (gray).

and seek to extract information encoded within it. UMAP permits customization of the distance metric used for embedding, so we could directly embed the data into target dimension 2 with toroidal topology (as demonstrated as an example in Ref. 4). However, the coordinates of this embedding did not generally match the true trajectory (Supp. Fig. 1F).

##### 1.3 Methods

We generate grid modules with different lattice angles as described in Methods of the main text (Sec. 4.1.2). Here, the transformation matrix from topological space to physical space used to generate tuning curves is

$$\mathsf{A} = l \begin{pmatrix} \cos\phi & \cos(\phi+\psi) \\ \sin\phi & \sin(\phi+\psi) \end{pmatrix}, \tag{1}$$

where $\psi$ is the lattice angle. As in the main text, we use grid scale $l = 40\,\mathrm{cm}$ and orientation $\phi = 0$. The animal trajectory is the same as that described in Methods of the main text (Sec. 4.1.1). The animal explores a 1.8 m-diameter circular environment for 1000 s, with timepoints sampled at 0.2 s intervals. The results in Supp. Fig. 1 are obtained using datasets consisting of 40 simulated grid cells from the same module.

We perform the same preprocessing steps of normalization, exclusion of silent timepoints, and geometric subsampling as done for persistent cohomology and described in Methods of the main text (Sec. 4.3). Subsampling selects 1000 out of 5000 timepoints as input into the algorithms for manifold learning. The learned transformation is then applied to all timepoints.

We implement Isomap with 5, 15, or 50 nearest neighbors [1]. The embeddings shown in Supp. Fig. 1A,B,D were obtained with 15 nearest neighbors. To obtain the visualization in Supp. Fig. 1A, we set the target dimension to 3. Otherwise, we set the target dimension to 4 and perform ICA decomposition on the Isomap embedding with the FastICA algorithm [2]. We test a variety of choices for the contrast function, including log-cosh, Gaussian, and quartic functions; the results in Supp. Fig. 1A–D were obtained with the log-cosh contrast function. Similar results (not shown) were also obtained using the FastISA algorithm [3] for Independent Subspace Analysis (ISA) with subspace dimension 2.

For each Isomap embedding, we run FastICA 3 times with random initial conditions. We partition each ICA decomposition with 4 components into pairs of components in all $\binom{4}{2} = 3$ ways. For each pair of components $(x_t, y_t)$ over timepoints $t = 1, \ldots, T$, we first calculate the normalized distance to the origin for each timestep:

$$r_t = \frac{\sqrt{x_t^2 + y_t^2}}{\frac{1}{T}\sum_t \sqrt{x_t^2 + y_t^2}}. \tag{2}$$

For a circular distribution, these distances $r_t$ should be lie near 1, so we define the *non-circularity* of the projection to be

$$\text{non-circularity} = \frac{1}{T}\sum_t (r_t - 1)^2. \tag{3}$$

Out of the 3 FastICA trials with 3 different partitions each, we select the partition whose component pairs have the lowest total non-circularity. Thus, we ultimately transform the Isomap embedding into 2 pairs of ICA components in which circular features are emphasized. For each component pair, we assign a circular coordinate to each timepoint vector by calculating its vector angle with respect to an arbitrary reference, which we take to be one coordinate axis in the pair. Thus, we obtain two circular coordinates for each timepoint in the Isomap embedding.

We implement UMAP with 5, 15, or 50 nearest neighbors and minimum distances of 0.1, 0.3, and 0.5 [4]. We obtained similar results across this range of parameters, and the embeddings shown in Supp. Fig. 1E,F were obtained with 15 nearest neighbors and a minimum distance of 0.3. To obtain the visualization in Supp. Fig. 1E, we set the target dimension to 3. Otherwise, we set the target dimension to 2 and define a toroidal metric for the embedding [4]. It has domain $[0, 2\pi) \times [0, 2\pi)$ and the distance between two points $(u_1, v_1)$ and $(u_2, v_2)$ is

$$d(u_1, v_1; u_2, v_2) = \sqrt{\langle u_1 - u_2 \rangle_\pi^2 + \langle v_1 - v_2 \rangle_\pi^2} \quad \text{where} \quad \langle a \rangle_\pi \equiv (a + \pi \mod 2\pi) - \pi \tag{4}$$

We take the coordinates of this embedding to be our circular coordinates directly.

To reconstruct the animal trajectory from circular coordinates, we proceed as in Methods of the main text (Sec. 4.5) for lattice angle 60°. For lattice angle 90°, we no longer need the unshearing steps, but the rest of the method proceeds accordingly.

### 2 Mathematical details for one-dimensional persistent cohomology

#### 2.1 Simplicial complexes

Given a set $P$, a *simplex* $\sigma$ is its subset, $\sigma \subseteq P$. The *dimension* of the simplex is one less than its cardinality, $\dim \sigma = \operatorname{card} \sigma - 1$. 0-, 1-, and 2-dimensional simplices are called vertices, edges, and triangles. If one simplex is a subset of another, $\sigma \subseteq \tau$, then $\sigma$ is called a *face* of $\tau$, and $\tau$ is a *coface* of $\sigma$. A simplicial complex $K$ is a collection of simplices closed under the face relation, i.e., for every simplex $\tau \in K$ and every $\sigma \subseteq \tau$, $\sigma \in K$.

Throughout the paper we use Vietoris–Rips complexes, which consist of all cliques in a nearest neighbor graph of a point set. Specifically, given a point set $P \in \mathbb{R}^n$ and a distance threshold $r$, the Vietoris–Rips complex consists of all subsets $\sigma$ of $P$, where every pair of points in the subset is within distance $r$ of each other,

$$\mathrm{VR}_r(P) = \{\sigma \subseteq P \mid \forall x, y \in \sigma, \|x - y\| \leq r\}.$$

#### 2.2 Cohomology

Let $\mathbb{Z}_p$ be the field of integers modulo a prime $p$. Given a simplicial complex $K$, let $K^0$, $K^1$, and $K^2$ denote its vertices, edges, and triangles. We define 0-, 1-, and 2-cochains as the functions from vertices, edges, and triangles into $\mathbb{Z}_p$.

$$\mathsf{C}^0(K; \mathbb{Z}_p) = \{\text{functions } f_0 : K^0 \to \mathbb{Z}_p\}$$
$$\mathsf{C}^1(K; \mathbb{Z}_p) = \{\text{functions } f_1 : K^1 \to \mathbb{Z}_p\}$$
$$\mathsf{C}^2(K; \mathbb{Z}_p) = \{\text{functions } f_2 : K^2 \to \mathbb{Z}_p\}$$

All of our results were obtained using $p = 3$.

Two *coboundary maps*, $d_0 : \mathsf{C}^0 \to \mathsf{C}^1$ and $d_1 : \mathsf{C}^1 \to \mathsf{C}^2$, are important for the construction of 1-dimensional cohomology used in the paper,

$$(d_0 f_0)(\{a,b\}) = f_0(b) - f_0(a)$$
$$(d_1 f_1)(\{a,b,c\}) = f_1(\{b,c\}) - f_1(\{a,c\}) + f_1(\{a,b\})$$

If for a function $f_1 \in \mathsf{C}^1$, $d_1 f_1 = 0$, then $f_1$ is a *cocycle*. If there is a map $f_0$ such that $d_0 f_0 = f_1$, then $f_1$ is a *coboundary*.

A fundamental property of the coboundary maps is that $d_1 d_0 f_0 = 0$. In other words, all coboundaries are cocycles, $\mathrm{im}(d_0) \subseteq \ker(d_1)$. The 1-dimensional cohomology is the quotient of the two groups,

$$\mathsf{H}^1(K; \mathbb{Z}_p) = \ker(d_1) / \mathrm{im}(d_0)$$

In other words, when two cocycles differ by a coboundary, we consider them equivalent, and the cohomology group consists of all equivalence classes of cocycles, called cohomology classes.

#### 2.3 Persistent cohomology

When we increase the distance threshold in the definition of the Vietoris–Rips complex, we get a larger nearest neighbor graph and therefore a larger set of cliques,

$$\mathrm{VR}_r(P) \subseteq \mathrm{VR}_s(P), \text{when } r \leq s.$$

When two complexes nest, there is a map on their cohomology groups, defined by restriction, $i : \mathsf{H}^1(L) \leftarrow \mathsf{H}^1(K)$, with $f_1^L = i(f_1^K)$ defined by $f_1^L(\{a,b\}) = f_1^K(\{a,b\})$ for every edge $\{a,b\} \in L$.

As we sweep the distance threshold $r$ from 0 to $\infty$, we get a nested sequence of Vietoris–Rips complexes, called a *filtration*. Passing to cohomology, we get a sequence of groups connected with the restriction maps,

$$\mathsf{H}^1(\mathrm{VR}_{r_1}(P); \mathbb{Z}_p) \xleftarrow{i_1} \mathsf{H}^1(\mathrm{VR}_{r_2}(P); \mathbb{Z}_p) \xleftarrow{i_2} \mathsf{H}^1(\mathrm{VR}_{r_3}(P); \mathbb{Z}_p) \xleftarrow{i_3} \ldots \leftarrow \mathsf{H}^1(\mathrm{VR}_\infty(P); \mathbb{Z}_p)$$

The theory of persistent (co)homology describes how to track cohomology classes as they appear and disappear in this sequence. If there is a class $\gamma$ that appears in $\mathsf{H}^1(\mathrm{VR}_{r_b})$, i.e., $\gamma \notin \mathrm{im}(i_b)$, and disappears by becoming a coboundary in $\mathsf{H}^1(\mathrm{VR}_{r_d})$, i.e., $(i_{d+2} \circ \ldots \circ i_{b-1})(\gamma) \in \ker(i_{d+1})$, then we have a birth–death pair $(r_b, r_d)$. The collection of all such pairs is called a *persistence diagram*.

#### 2.4 Circular coordinates

Replacing integers modulo $p$, $\mathbb{Z}_p$, with integers, $\mathbb{Z}$, in the definition of the cohomology group, we get a cohomology group with integer coefficients, $\mathsf{H}^1(K; \mathbb{Z})$. Every cocycle $f_1$ in this group can be interpreted as a continuous map from the complex $K$ into the circle, where all vertices map to the same point on the circle, and every edge $\{a,b\}$ winds around the circle the number of times prescribed by the integer $f_1(\{a,b\})$, with positive integers giving a clockwise and negative integers counter-clockwise winding.

de Silva et al. [6] describe a procedure that selects a persistent cocycle, lifts it to a cocycle with integer coefficients, reinterprets the latter as a map to a circle, and then continuously smooths the vertices around the circle via least squares optimization, resulting in a map from the input point set to the circle that captures the underlying topology of the data.

Perea [7] extends the circular coordinates construction to work with a simplicial complex computed on a subsample of the data (e.g., the geometric subsample described in the paper) to the full data set. This extension grants a significant computational advantage.

##### 2.5 Software for circular coordinates

The algorithm of de Silva et al. [6] is available as part of Dionysus[1]. The algorithm of Perea [7] is implemented in DREiMac[2], which uses Ripser[3] to compute persistent cohomology. We use Ripser and DREiMac throughout the paper.

[3] Aapo Hyvärinen and Urs Köster. FastISA: A fast fixed-point algorithm for independent subspace analysis. In *ESANN2006: 14th European Symposium on Artificial Neural Networks*, 2006.

[4] Leland McInnes, John Healy, Nathaniel Saul, and Lukas Großberger. UMAP: Uniform Manifold Approximation and Projection. *Journal of Open Source Software*, 3(29):861, 2018.

[5] Benjamin Dunn. Toroidal topology of grid cell ensemble activity. International Conference on Mathematical Neuroscience, 2020.

[6] Vin de Silva, Dmitriy Morozov, and Mikael Vejdemo-Johansson. Persistent cohomology and circular coordinates. *Discrete & computational geometry*, 45(4):737–759, June 2011.

[7] Jose A Perea. Sparse circular coordinates via principal $\mathbb{Z}$-Bundles. *arXiv*, September 2018.

[1] `github.com/mrzv/dionysus`
[2] `github.com/ctralie/DREiMac`
[3] `github.com/ripser/ripser`
